# Acute and Chronic Molecular Signatures and Associated Symptoms of Blast Exposure in Military Breachers

**DOI:** 10.1101/738054

**Authors:** Zhaoyu Wang, Caroline M Wilson, Natalia Mendelev, Yongchao Ge, Hanga Galfalvy, Gregory Elder, Stephen Ahlers, Angela M Yarnell, Matthew L LoPresti, Gary Kamimori, Walter Carr, Fatemeh Haghighi

## Abstract

Injuries from exposure to explosions rose dramatically during the Iraq and Afghanistan wars, which motivated investigations of blast-related neurotrauma and operational breaching. In this study, military “breachers” were exposed to controlled, low-level blast during a 10-day explosive breaching course. Using an omics approach, we assessed epigenetic, transcriptional, and inflammatory profile changes in blood from operational breaching trainees, with varying levels of lifetime blast exposure, along with daily self-reported symptoms (with tinnitus, headaches, and sleep disturbances as the most frequently reported). Although acute exposure to blast did not confer epigenetic changes, specifically in DNA methylation, differentially methylated regions (DMRs) with coordinated gene expression changes associated with chronic lifetime cumulative blast exposures were identified. The accumulative effect of blast showed increased methylation of PAX8 antisense transcript with coordinated repression of gene expression, which has been associated with sleep disturbance. DNA methylation analyses conducted in conjunction with reported symptoms of tinnitus in the low vs. high blast incidents groups identified DMRS in KCNE1 and CYP2E1 genes. KCNE1 and CYP2E1 showed the expected inverse correlation between DNA methylation and gene expression, which have been previously implicated in noise related hearing loss. Although no significant transcriptional changes were observed in samples obtained at the onset of the training course relative to chronic cumulative blast, we identified a large number of transcriptional perturbations acutely pre- versus post-blast exposure. Acutely, 67 robustly differentially expressed genes (fold change ≥1.5), including UFC1 and YOD1, ubiquitin-related proteins were identified. Inflammatory analyses of cytokines and chemokines revealed dysregulation of MCP-1, GCSF, HGF, MCSF, and RANTES acutely following blast exposure. These data show the importance of an omics approach, revealing that transcriptional and inflammatory biomarkers capture acute low-level blast overpressure exposure, whereas DNA methylation marks encapsulate chronic long-term symptoms.

## INTRODUCTION

Injuries from exposure to explosive blasts rose dramatically during Operation Iraqi Freedom and Operation Enduring Freedom (OIF, OEF) due to increased use of improvised explosive devices (IEDs) in military settings and has also increased in civilian populations through acts of terrorism^1-3^, thus motivating investigations of blast-related neurotrauma. Congressional acts including the John S. McCain National Defense Authorization Act for Fiscal Year 2019 have called for “review of guidance on blast exposure during training” (Section 253), which emphasize the importance of understanding and mitigating the effects of blast after successive days of training and over the course of a military career.^4^ Despite the increase in occurrence and public awareness, our understanding of the effects of blast and the mechanisms behind subsequent injury are limited.^5, 6^ Blast overpressure (BOP) events are capable of injuring the brain by means of high energy pressure waves that are rapidly emitted from the explosive, which propagate from the object and turn into shock waves upon interacting with a medium—in this case being the military warfighter.^7^ The clinical and pathological effects of blast exposure varies depending on the magnitude of the detonation, proximity to the blast, and the use of protective gear^7–9^, and includes but is not limited to neuronal swelling, subdural hematomas, myelin deformation, inflammation, loss of consciousness, temporary disorientation, sleep disturbances, memory deficits, and tinnitus.^10–18^ A better understanding of the systemic biomarkers underlying blast exposure responses is critical to not only the identification of blast-related injury, but also to the development of effective diagnostics and potential treatments.

In this effort, we have undertaken prospective human studies involving “Breachers,” military and law enforcement personnel who are exposed to repeated blast as part of their occupational duty. Breachers are typically in close proximity to controlled, repeated, low-level blast during operations and training, and have reported a range of physical, emotional, and cognitive symptoms, including headache, sleep disturbance, anxiety, and impaired cognitive performance.^19^ It has been previously shown that blast overpressure exposure is capable of inducing changes in gene expression in military personnel^20^ and in animal models of blast injury.^21^

We took an omics approach, performing large-scale epigenetics, as well as transcriptional, and inflammatory profiling to identify blood-based biomarkers associated with acute and chronic blast exposure. We investigated DNA methylation, a highly stable epigenetic marker associated with gene repression as well as gene expression patterns, in addition to transcriptome analysis via RNA-seq to identify potential epigenetic and coordinated gene expression abnormalities. Furthermore, we assessed inflammatory changes following blast exposure, in addition to self-reported symptoms. These data allow us to investigate regulatory, transcriptional, and inflammatory biosignatures of blast exposure.

## METHODS

### Samples & Subjects

This study was approved by the Institutional Review Board (NMRC#2011.0002; WRAIR#1796). The present study obtained samples from 34 healthy, male participants at U.S. Army explosive entry training sites (special operations and combat engineer courses). Biological specimens for epigenetics and transcriptional studies were collected at the start (baseline) and at the end of the training course (pre- vs post-blast exposure), and serum samples for inflammatory analyses were collected daily. Demographic information including sex, age, lifetime operational exposure to blast, as well as self-reported TBI history were recorded at the start of the training. During the course of the training, self-report symptom assessments were completed daily.

### DNA methylation sample processing & Quality Control

Blood was collected using EDTA tubes, and processed to separate peripheral blood mononuclear cells (PBMCs). PBMCs were purified by Ficoll gradient, washed with PBS and stored at −80°C. DNA was extracted with QIAamp DNA Micro Kit (QIAGEN, Hilden, Germany). Genomic DNA was bisulfite converted (Zymo Research, Irvine, CA, USA) and CpG methylation was determined using Illumina Infinium HumanMethylationBeadChip microarrays (HM450, Illumina, Inc., San Diego, CA, USA), as described previously.^22^ Data & quality control (QC) analyses were performed using R Language 3.03 ^23^, an environment for statistical computing, and Bioconductor 2.13.^24^ Raw data files (.idat) were processed by minfi package.^25^ All samples displayed a mean probe-wise detection call for the 485512 array probes < 0.0005 (Figure S1). Sex QC analysis, also confirmed methylation-based sex prediction with those reported (Figure S2). For QC sample tracking of pre- vs. post-Breacher training, we used the 65 single nucleotide polymorphism (SNP) probes included in the HumanMethylation 450k BeadChip, confirming that subjects for which multiple samples were available grouped together (Figure S3).

### DNA methylation data analysis

For all DNA methylation analyses, we used the matrix of M-values (logit transformation of beta-values) which correspond to methylation levels. Surrogate variable analysis (SVA) was performed to add surrogate variables and rule out potential batch effects. A linear model was used for the binary variable of interest, while including age and history of TBI as covariates in the model. Performing the comparative analysis in limma^26^ implemented in R, we obtained t-statistics and associated p-values for each CpG site. The point-wise p-values, were then used for the identification of differentially methylated regions (DMRs) using the combined-p-value tool.^27^

### Total RNA sample/ library preparation and sequencing

Blood was collected using Paxgene RNA tubes (PreAnalytiX, QIAGEN/BD, Hombrechtikon, Switzerland) according to manufacturer’s instructions and stored at −80°C. RNA was extracted with the Paxgene Blood RNA Kit (PreAnalytiX). Globin mRNA was removed with Globin Clear Human Globin mRNA Removal kit (Ambion, Inc., Austin, TX, USA). All RNA samples had RNA integrity numbers (RIN) ≥6.0. Total RNA sequencing libraries were prepared using the Illumina Stranded Total RNA Library Prep Kit with Ribo-Zero Gold (Illumina, Inc.) in accordance with the manufacturer’s instructions. Briefly, 290ng-500ng of total RNA was used for ribosomal depletion and fragmented by divalent cations under elevated temperatures. The fragmented RNA underwent first strand synthesis using reverse transcriptase and random primers followed by second strand synthesis to generate cDNA. The cDNA fragments underwent end repair, adenylation and ligation of Illumina sequencing adapters. The cDNA library was enriched using 11 cycles of PCR and purified. Final libraries were evaluated using PicoGreen (Life Technologies, Carlsbad, CA, USA) and Fragment Analyzer (Advanced Analytical, Agilent Technologies, Santa Clara, CA, USA) and were sequenced on an Illumina HiSeq2500 sequencer (v4 chemistry) using 2 x 125bp read lengths.

### RNA-seq data preprocessing and bioinformatics analysis

Reads were aligned to the human reference hg19 using STAR aligner (v2.4.0c).^28^ Quantification of genes annotated in Gencode v18 was performed using featureCounts (v1.4.3). QC metrics were collected with Picard (v1.83) and RSeQC ^29^ (http://broadinstitute.github.io/picard/). Normalization of feature counts was done. Furthermore, for gene expression analysis, we used the voom function in limma to get logCPM matrix, where the design matrix consists of intercept, age, history of TBI, and variable of interest. We used SVA to rule out potential batch effects. Analyses were performed in limma, and point-wise as well as multiple testing adjusted p-values were reported.

### Gene Ontology & Gene Set Enrichment Analyses

We performed Gene Ontology (GO) analysis using goseq R package, with gene length bias considered. Gene set enrichment analysis (GSEA) version 3.0 ^30^ was run on our ranked list of 8,157 genes, which is filtered from the logCPM matrix obtained in pre-post analysis by criteria that average logCPM is greater than or equal to 4 in either of the comparison groups considered here, and ordered by t-statistics. GSEA preranked was run with 1000 permutations using gene sets from the Molecular Signatures Database (MsigDB) ^31^ as follows: 1) gene ontology gene sets (C5) including biological processes (BP), cellular components (CC), and molecular function (MF); and 2) hallmark gene set (H).

### Cytokine sample processing via Luminex

For assays of inflammatory cytokines, we used the Luminex 63-Plex assay (eBiosciences/Affymetrix, Inc., Santa Clara, CA, USA), which has the ability to multiplex, measuring levels of 63 inflammatory cytokines simultaneously. Assays were performed by the Human Immune Monitoring Center at Stanford University and kits were used according to the manufacturer’s recommendations with modifications as described here briefly. Beads were added to a 96 well plate and washed in a BioTek Elx405 washer (BioTek Instruments, Inc., Winooski, VT, USA). Serum samples were added to the plate containing the mixed antibody-linked beads and incubated at room temperature for 1 hour followed by overnight incubation at 4°C with shaking. Cold and room temperature incubation steps were performed on an orbital shaker at 500-600 rpm. Following the overnight incubation, plates were washed in a BioTek Elx405 washer, and then biotinylated detection antibody was added for 75 minutes at room temperature with shaking. Plates were washed as above and streptavidin-PE was added. After incubation for 30 minutes at room temperature, a wash was performed as above and reading buffer was added to the wells. Each sample was measured in duplicate. Plates were read using a Luminex 200 instrument (Millipore Sigma, Burlington, MA, USA) with a lower bound of 50 beads per sample per cytokine. Custom assay control beads by Radix Biosolutions (Georgetown, TX, USA) were added to all wells.

### Cytokine data analysis

Cytokine data was available for a subset of subjects, for 5 days: Day 4, Day 7, Day 8, Day 9 and Day 10. Two levels were taken for each subject for each day, and all values were analyzed. All cytokine levels were log-transformed, and then graphed. Since outliers were present in many of the cytokines, values were capped (winsorized) from both below and above, at the lower quartile minus 1.5 times the Inter Quartile Range (IQR), and the upper quartile plus 1.5 times the IQR, respectively. Separate mixed effect models were fit for each log-transformed and capped cytokine measure, with day as fixed, categorical predictor, and subject-specific random intercepts. P-values for each level of the fixed effect were recorded, and corrected for multiple testing (for K=63 cytokines) using the Bonferroni method. To calculate standardized effect sizes for the contrast of each experimental day compared to the baseline, the model coefficients were divided by the standard deviation of the baseline measure, with the scaleless effect sizes then represented via heatmap plots.

## RESULTS

The present study collected data from 34 participants during 3 separate 2-week data collection cycles at U.S. Army explosive entry training sites (special operations and combat engineer courses). In these advanced training courses, both trainees and instructors have a career history of repeated exposure to low level blasts. Blood samples were obtained pre- and post-training for epigenetics, transcriptional, and protein assays, subsequently referred to as pre-vs post-blast exposure. All participants were male, with an average age of 30.79 years (S.D. 4.57 years). Self-report history of injury and of blast exposure was recorded at baseline, and daily self-report symptom assessment during the training course was also recorded (Supplemental Figure S4). The chronology of exposures during the 2-week Breacher training and participants’ reported history of lifetime exposure to blast and TBI history is provided in Figure 1A & B, demonstrating participant exposures ranging from tens to hundreds over a military career. A total of 60% of these 34 breachers self-reported at least one lifetime mild traumatic brain injury (TBI) event; however, there was no correlation between history of TBI and the number of lifetime blast exposures (p > 0.8).

**Figure 1:**
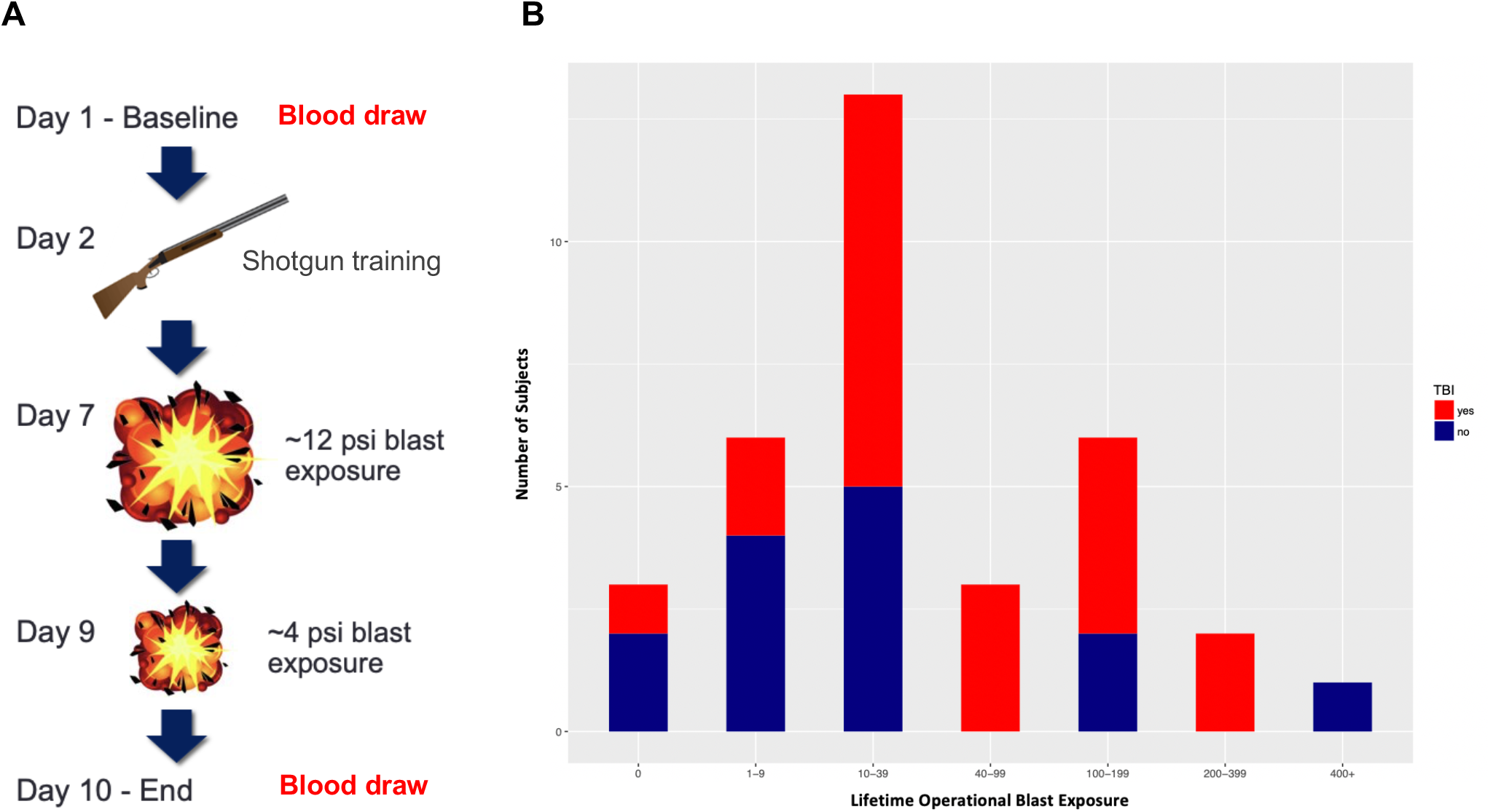
(A) Protocol for operational breaching and blast exposure throughout the 10-day training course, showing pre- and post-blast exposure blood draws on Days 1 and 10, respectively. (B) Distribution of number of lifetime operational blast exposures with history of self-reported TBI (red) and no history of TBI (blue).

### DNA methylation & transcriptional changes associated with chronic cumulative career blast exposures

For baseline low vs. high career blast exposure DNA methylation analyses, we used 34 subjects for which baseline data were available to examine whether the number of cumulative blast exposure events during a career in military service was associated with changes in transcriptional regulation. We empirically defined the low exposure group as those with less than 40 reported blast exposures and the high group reporting greater numbers. We asked whether the number of cumulative blast exposure events during a career in military service is associated with changes in transcriptional regulation. Whole genome transcriptional profiling via RNA-seq did not show significant gene expression changes between low vs. high blast lifetime exposed groups following multiple testing correction (data not shown). However, DNA methylation analyses of these samples via Illumina 450K methylation microarrays identified significant methylation differences. We found 10 significantly differentially methylated regions (DMRs) and genes associated with cumulative blast (Figure 2A and Supplementary Table S1). The majority of DMRs exhibited gain of methylation associated with cumulative blast exposure, with corresponding gene expression changes (Figure 2B). The PAX8 gene is an antisense transcript within the PAX8 gene, wherein the DMR is localized in promoter of this antisense transcript (Figure 2C). Interestingly, gain of DNA methylation in those subjects with high cumulative blast exposure corresponds to loss of gene expression in the PAX8 antisense transcript (NR_047570) (Figure 2B-C). PAX8 transcription factor is responsible for control of expression of thyroid specific genes involved in thyroid function, and, more recently, genome-wide association studies have implicated variants associated with PAX8 and sleep duration.^32–34^

**Figure 2:**
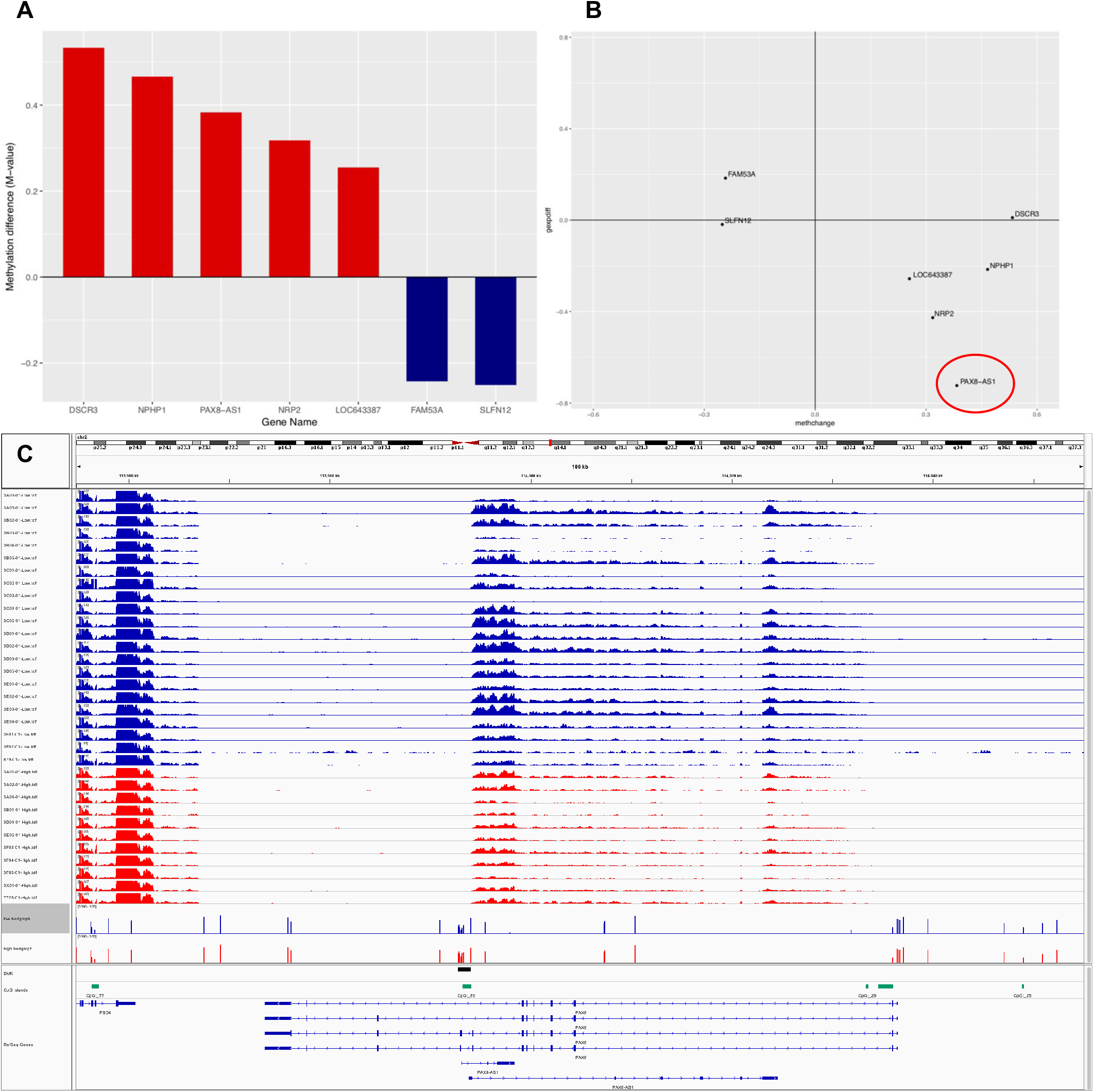
Differentially methylated regions (DMRs) associated with accumulative blast exposure. (A) DMRs with corresponding observed methylation data, showing gain (red) and loss (blue) of DNA methylation in low vs. high lifetime exposure to blast events. (B) DNA methylation and gene expression changes, with the majority of genes showing the expected anti-correlation between DNA methylation and gene expression. (C) Genome browser representation of PAX8 antisense transcript expression relative to accumulative lifetime blast exposure in low (blue) vs high (red) exposed breacher groups. Following an anticorrelated pattern of gain of DNA methylation and repression of gene expression, we see that the low accumulative blast exposed group shows a loss of DNA methylation DMR track in blue, whereas the high accumulative blast exposed group shows a gain of methylation indicated in red in the DMR track with coordinated repression of gene expression, as shown per subject. It should be noted that the DMR seems to be specifically regulating the PAX8 antisense transcript since the upstream gene PSD4 does not seem to be impacted by the differential methylation at the DMR.

### DNA methylation & transcriptional changes associated with acute blast exposure

31 of the subjects had available samples for both the baseline timepoint and at day 10 (completion of Breacher training). Comparing DNA methylation patterns pre- vs. post-blast exposure, no differences were identified. Interestingly, however, gene expression analyses of RNA-seq data revealed 6362 genes with statistically significant differential expression following multiple testing corrections (Figure 3A and Supplementary Table S2). To investigate robust gene expression changes associated with acute blast exposure, we focused on those genes with moderate-to-large fold change in expression (≥1.5 FC) and at least moderate expression levels in pre- or post-assessments (logCPM≥4). Using these stringent criteria, we identified 67 genes that show robust fold-change pre- vs. post-blast exposure (Figure 3B). An overwhelming number of these genes are involved in ribosomal functioning, which were dysregulated following blast. Transcripts of ribosomal proteins constituted 30% of these genes, with additional related transcripts of chaperone proteins involved in protein biogenesis and degradation (i.e., heatshock protein HSP90, translation elongation factors EEF1B2, EEF1A1P5, EEF1A1, ribosome biogenesis homolog [NSA2], and ubiquitins UFC1 & YOD1). Some of these loci and genes in related pathways have been previously implicated in responses to cellular stress, neurodegeneration, and TBI.^35–38^ Of note, ubiquitin proteins have been implicated in TBI and neurodegenerative disease, and the ubiquitin C-terminal hydrolase-L1 (UCH-L1) has been FDA approved as a robust blood-based biomarker for acute mild brain injury.^39^

**Figure 3:**
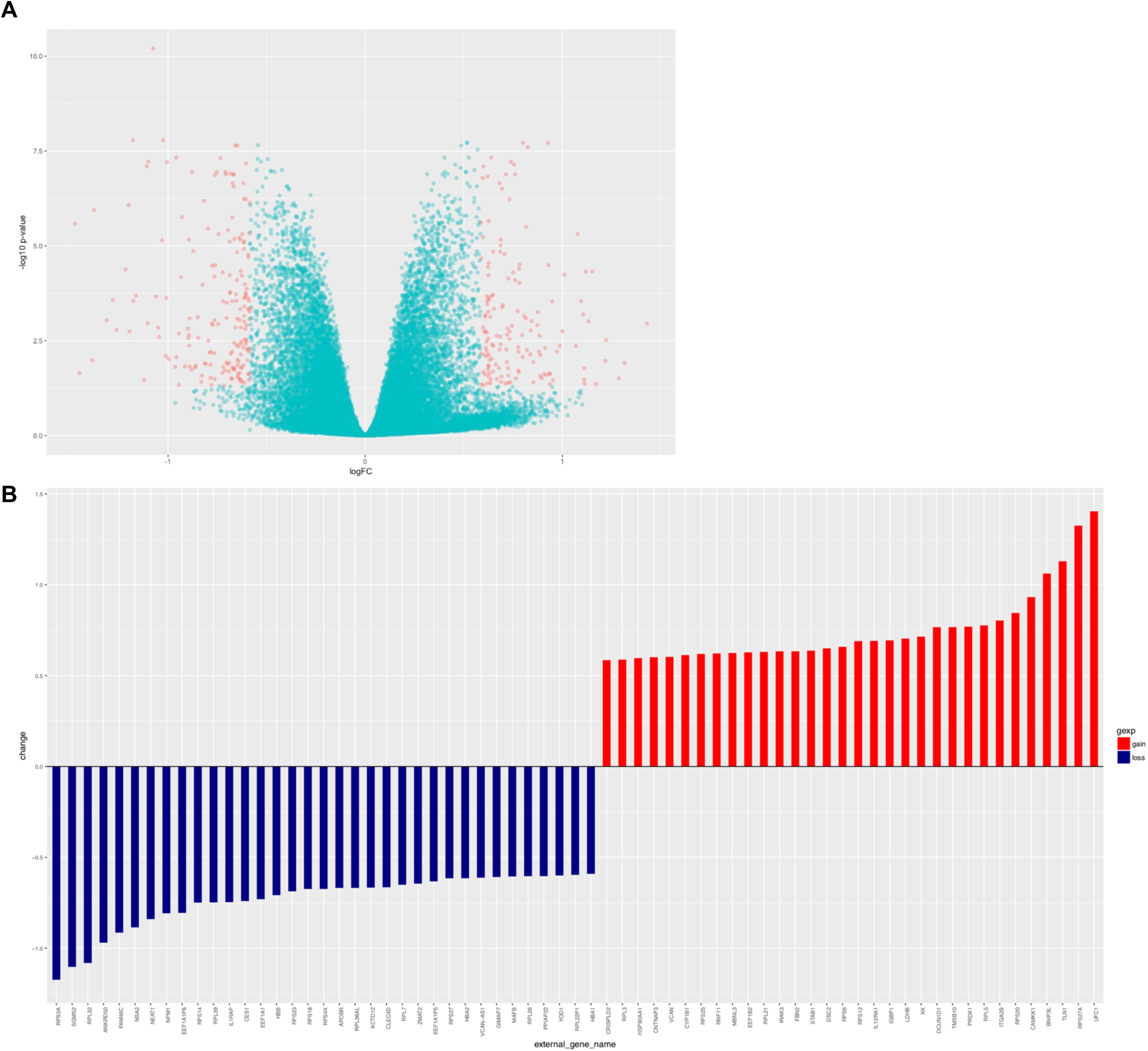
Gene expression changes tracks molecular response to acute exposure to blast. (A) Volcano plot highlighting genes that passed multiple testing correction (6362 genes), that also showed greater than ±1.5 fold change in expression (336 genes, red). The y-axis shows the unadjusted p-value. (B) Showing genes (67) with robust expression changes pre vs. post blast training with fold change ≥ |1.5| and excluding rarely expressed genes (logCPM<4).

### Blast associated physiological and psychological symptoms linked to DNA methylation and transcriptional changes

Given that daily symptom reports were ascertained from all participants, we utilized this information to track DNA methylation and gene expression changes associated with symptom reporting. Given the data sparsity, with missing symptom reporting by some participants, we performed initial symptom filtering. We kept those symptoms which were endorsed by ≥10 subjects for each comparison. For the baseline analysis of the high vs low cumulative blast exposed groups (day 1 symptom report), we observed a higher average symptom score (>0.25) in the high vs. low lifetime blast groups and filtering produced one symptom, tinnitus (ringing in the ear), with 11 participants reporting (Figure 4A). Tinnitus is a commonly reported symptom by military and Veteran subjects with repeated exposure to blast.^19^ Although we found no significant genome-wide transcriptional changes following multiple testing correction, we report point-wise transcriptional changes for the most robustly differentially expressed loci (with FC ≥±1.5 Supplementary Table S3). We did identify genes with differential DNA methylation changes that track with reported symptoms of tinnitus in the high vs. low career breaching groups (Figure 4B). Of these differentially methylated regions and associated loci potassium voltage-gated channel, Isk-related family, member 1 (KCNE1), Cytochrome P450 family 2, subfamily E, member 1(CYP2E1), dual specificity phosphatase 22 (DUSP22), and hyperpolarization activated cyclic nucleotide gated potassium channel 2 (HCN2) genes have been previously implicated in auditory functioning.^40–42^ To determine whether these DMRs confer transcriptional regulatory changes, we examined the corresponding gene expression levels in the low vs. high breaching groups in these loci. We found that the genes KCNE1 and CYP2E1 showed the expected anticorrelated pattern of gain in DNA methylation and loss of gene expression (Figure 4C). KCNE1 is associated with noise related hearing loss through human genetic studies and animal studies have also implicated CYP2E1 as associated with nitrile exposure and noise related hearing loss in rodents.^43–45^

**Figure 4:**
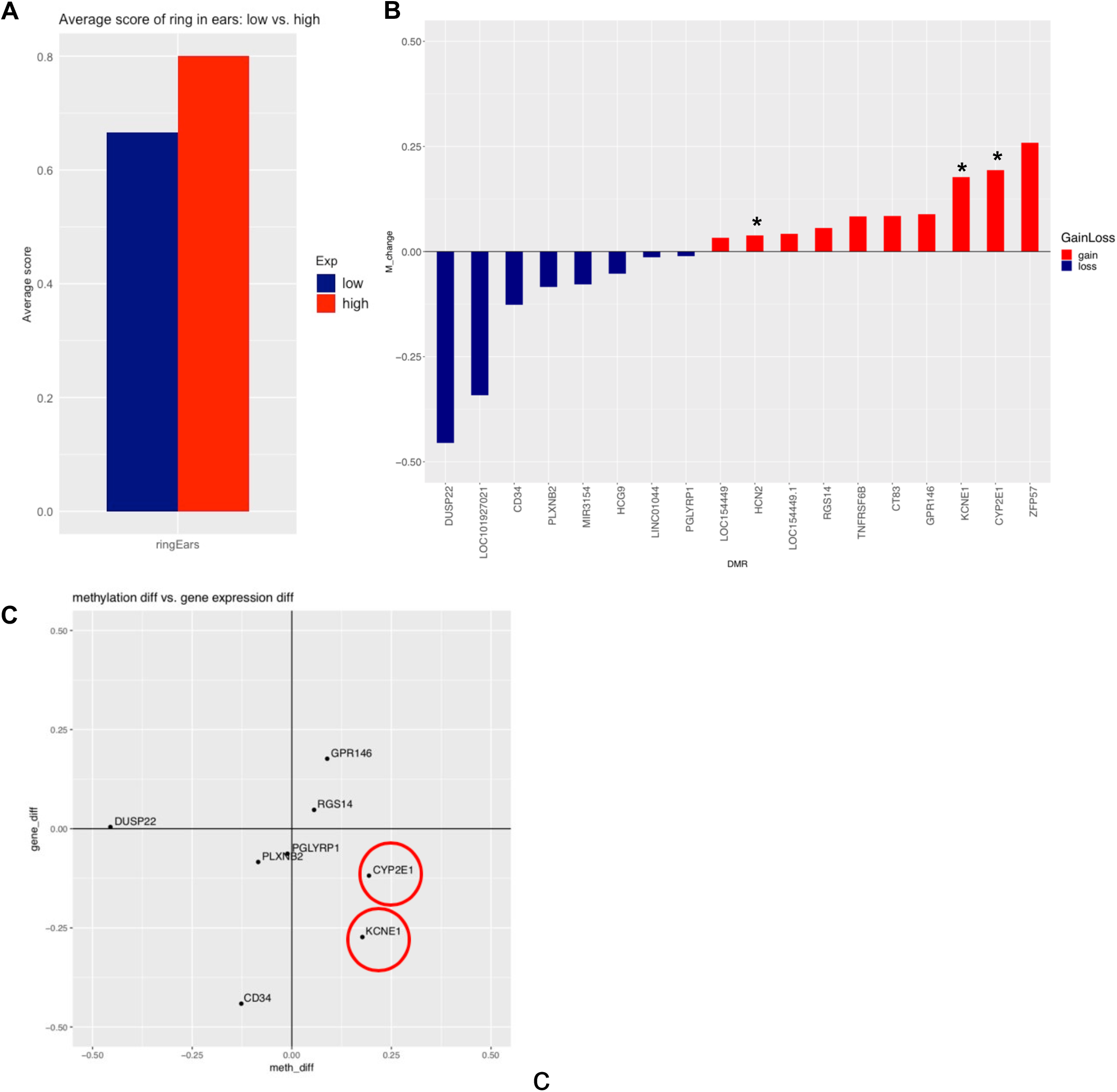
(A) To determine the effect of cumulative blast exposure on physiological and psychological symptoms, we examined average symptom levels at baseline for those with low vs high career breaching, with ringing in the ear being the top reported symptom. (B) Differentially methylated genes associated with ringing in the ear/tinnitus in low vs. high career breaching, denoting genes implicated in auditory functioning by (*). (C) Of note, KCNE1 and CYP2E1 genes follow the expected anti-correlated pattern of increase in DNA methylation with decrease in gene expression.

For the pre-post blast exposure analysis, we accounted for the observation that the breachers had a higher exposure to blast on day 7 (average exposure of ∼12psi; Figure 1A), well beyond the 4psi exposure level used as a safety threshold on most training ranges.^46^ Typical exposures in this type of training are less than 1psi.^47^ For consideration of the large magnitude blast exposure we recorded, we took the average symptom levels reported in days 1-6 and compared with the average from days 7-10, inclusive of the 7th day high magnitude blast event. Similarly, for pre-post breaching course analyses, we analyzed symptoms that ≥10 subjects had when responding pre- vs. post-blast, where pre corresponds to symptoms reported at least once during training days 2-6 and post days 7-10, with higher (again >0.25) average symptom scores in post- vs. pre-days. We chose this separation interval, because the breaching cohort was exposed to a higher level of blast (max ∼12psi) during day 7 of training. Using an *a priori* sample size criterion of ≥10 participants reporting, we found that headache was the most highly reported symptom pre-post blast exposure, endorsed by 18 subjects (Figure S4). In line with the pre-post DNA methylation results on the total subjects, the symptom analysis also did not show statistically significant genome-wide DMRs that tracked with reported symptoms of headache post blast exposure, following multiple testing correction. Also, no significant transcriptional changes were detected with multiple testing correction, yet pointwise data for the most significant gene expression changes with ≥±1.5 fold change are reported in supplementary data (Table S4).

### Inflammatory markers of acute blast exposure

Assaying cytokine levels across the training course allows us to determine how repeated exposure to blast and breaching environment induces perturbations in levels of cytokine proteins. Data was available for 32 subjects. Cytokine measures on day 4 were used as a reference, because by this time the participants had acclimated to the training environment and had no exposure to high explosive blast or physical exertions on that day. Comparatively, days 7-10 were considered post-blast exposure days respectively. We contrasted cytokine levels for each of the post-blast days (7-10) to those from the reference (day 4). Comparing cytokine levels pre-post exposure, we found significant elevation in five out of 63 cytokines (MCP1, GCSF, HGF, MCSF and RANTES) associated with acute blast exposure (Figure 5; p≤0.05 corrected for multiple testing). These data show that changes in cytokines track acute exposure to blast, as indicated by the observed effect sizes depicted in Figure 5 (with effect sizes for all cytokines shown in Supplemental Figure S5 and Supplementary Table S5).

**Figure 5:**
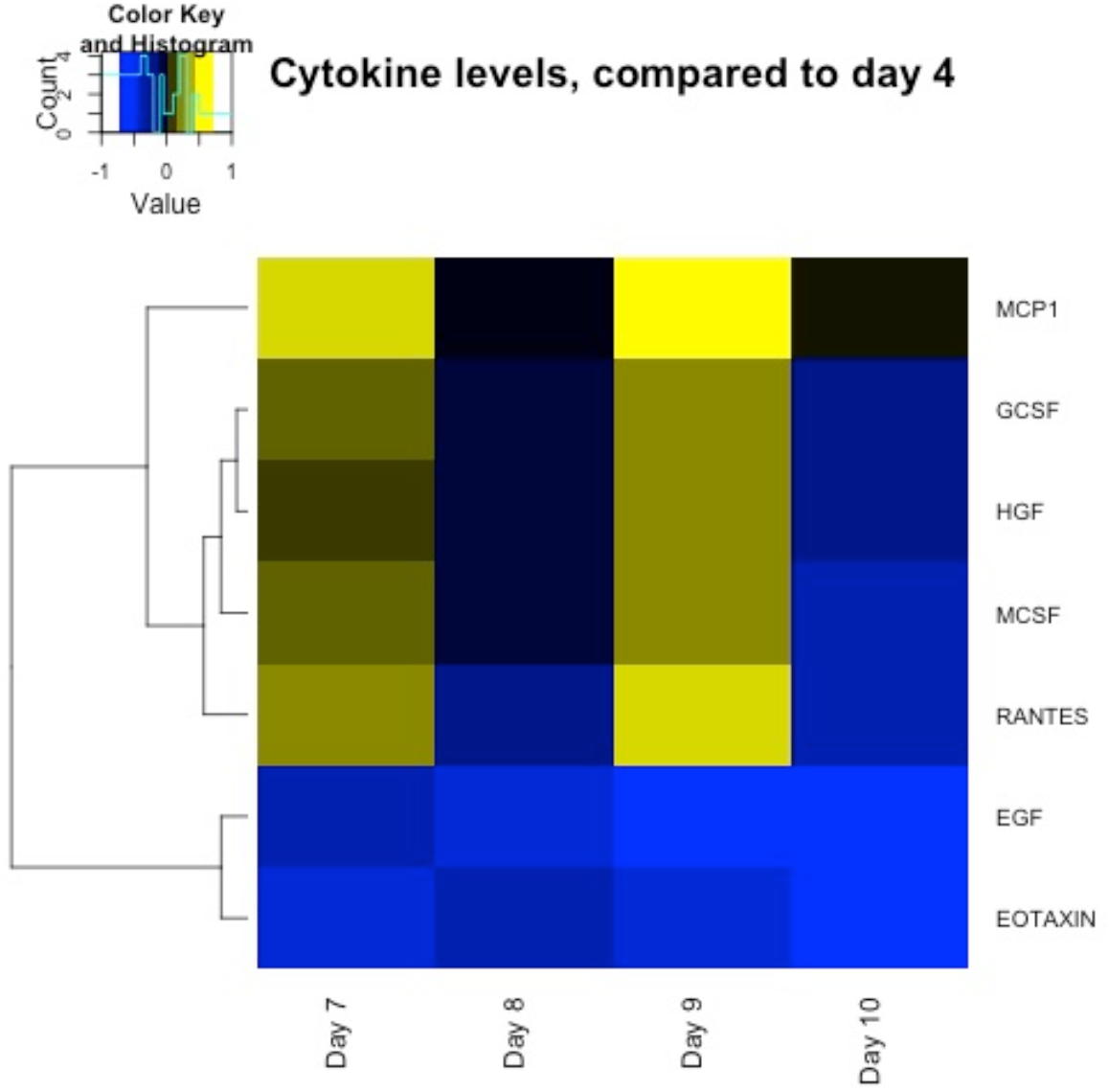
Heat map showing significant changes in cytokine level tracking with blast exposure (days 7 and 9) using day 4 (no exposure to blast activity) as the reference (Bonferroni corrected).

## DISCUSSION

In the present study, we have taken a multimodal omics approach to elucidate changes in inflammatory, epigenetic, and transcriptional profiles of military breachers following a 10-day breacher operations training course. We obtained DNA methylation, RNA-seq, and chemokine/cytokine data for military subjects before and after blast exposure in order to identify changes in DNA methylation, transcriptional, and inflammatory processes, which could serve as potential acute & chronic biomarkers of blast exposure in neurotrauma patients. Cumulative blast exposure history was used in conjunction with epigenetic and transcriptional data in order to determine the extent to which low or high numbers of exposure to repeated blast impacts physiological responsivity. Furthermore, self-reported symptom information was used in order to identify novel gene associations with blast-related symptomology.

A major limitation of the study is that DNA methylation and gene expression changes detected may not be reflective of acute changes associated with blast exposure, due to the blood collection end point on day 10. Although this likely has a bigger effect in regards to gene expression rather than DNA methylation, future studies would require specimen collection directly following blast exposure training sessions in order to more accurately assess transient changes in methylation and transcriptional processes. An additional limitation of the study is the lack of longitudinal data, for both genetic and symptom-based studies, on the breacher participants. Increased collection points several weeks or months after blast exposure training would allow for a better understanding of the dynamics and long term effects of blast exposure on the epigenetic, transcriptional, and symptom-based profiles in blast-exposed individuals. Furthermore, we acknowledge that our study lacks representation of both sexes, for the present cohort involved only male participants; thus findings may not be generalizable to females.

The primary findings of our study suggest that DNA methylation and gene expression are modulated by blast exposure in military personnel following blast exposure training programs. Although we were unable to detect acute changes in DNA methylation pre- vs. post-blast exposure, we show that high cumulative blast exposure history alters DNA methylation patterns relative to subjects with lower blast exposure throughout their military careers. Most notably, these DNA methylation changes associated with cumulative exposures generally conferred functional changes in gene expression. Specifically, increase in DNA methylation in the promoter region of the paired box gene 8, PAX8, antisense transcript in breachers with high cumulative blast exposure history (Figure 2A), conferred decrease in PAX8 gene expression (Figure 2B). This is potentially important mechanistically, because the PAX8 transcription factor is involved in the control of the expression of thyroid specific genes and thyroid development ^33, 48^, and it has recently been associated with alterations in sleep duration.^32, 34^ The association between PAX8 and sleep duration is particularly relevant, for sleep dysregulation is reported in our breacher symptom data (Figure S4) and in both active duty service members and US Veterans.^49, 50^ Reciprocally, sleep dysregulation has been shown to impact both metabolic and endocrine function^51^, possibly linking PAX8 with modulatory relationship between sleep dysregulation and endocrine processes.

This finding associating high lifetime cumulative blast exposure to changes in PAX8 antisense transcript methylation (and its respective association with changes in sleep duration) is important because sleep disturbances are reported in military warfighters in both active duty and post-deployment settings.^49, 50, 52^ Sleep disturbances lead to immediate impairments in cognition and alertness potentially resulting in accidents or death^53^, and are also associated with mental health outcomes including PTSD and depression in military populations.^49, 52^ While the changes in DNA methylation of PAX8 represent more stable marks as chronic biosignatures of lifetime blast exposure history.

Acute transcriptional alteration post-blast exposure in the KCTD12 gene was previously shown to be involved in circadian regulation ^54^, were also detected in this study. Potassium channel tetramerization domain containing protein (KCTD) 12, an auxiliary subunit of the GABA_B_ receptor gene, has been shown to both increase desensitization and slow the onset of GABA_B_ G protein-coupled receptor responses.^55, 56^ Previous animal studies by Cathomas et al. generated *Kctd12* null mutant (*Kctd12^-/-^*) and heterozygous (*Kctd12^+/-^*) mice in order to determine if behavioral and neurobiological observations could be made that are in line with the endophenotypes observed in psychiatric populations with KCTD12 perturbations.^54^ Notably, *Kctd*^-/-^ and *Kctd*^+/-^ mice show increased electrical excitability of CA1 pyramidal neurons ^54^, further implicating KCTD12 in mediating synaptic transmission and neuropsychiatric endophenotypes. Phenotypically, mice are typically inactive during the light phase of their circadian rhythm cycle ^57^, however, *Kctd12^+/-^* mice exhibit increased activity during the inactive (light) phase of the circadian cycle^54^, suggesting that downregulation of KCTD12 modulates circadian rhythm cycles. Interestingly, we also found KCTD12 as significantly downregulated pre-post blast exposure. Our RNA-seq findings in conjunction with our DNA methylation findings further associate the effects of blast overpressure exposure with biological/molecular perturbations potentially resulting in sleep dysregulation in our military warfighters.

Furthermore, RNA-seq data revealed 67 genes (32 upregulated; 35 downregulated) that had robust changes in gene expression (fold expression changes greater than or equal to 1.5) following exposure to blast (Figure 3B). We identified a subset of these 67 significantly differentially expressed genes that are involved in cell-cell adhesion, fibrosis, and accumulation of extracellular matrix (ECM) proteins. Specifically, VCAN encodes the proteoglycan versican which is one of the primary proteins that makes up the ECM.^58^ FBN2 regulates processes related to elastic fiber assembly and both TLN1 and DSC2 are implicated in cell-cell adhesion.^59–61^ It may be posited that increase expression of these genes as a protective mechanism following blast exposure in order to allow cells to potentially better resist the harmful effects of mechanical stress. We speculate that the notable increases in expression in CRISPLDL2 and STAB-1, genes involved in angiogenesis and lung morphogenesis, are likely modulated in response to blast exposure rather than related to respiratory processes. Decrease in expression of HBA1, HBA2, and HBB (hemoglobin subunit alpha 1, hemoglobin subunit alpha 2 and hemoglobin subunit beta, respectively [Figure 3B]), may suggest that the increase in expression of angiogenesis-related genes could be associated to the body’s injury response rather than physiological oxygen needs. These gene expression findings were corroborated by Gene Set Enrichment Analyses (GSEA), where positive enrichment in gene sets related to vasculature development, blood vessel morphogenesis, wound healing, and cortical cytoskeleton organization were also observed (Supplementary Table S6).

Furthermore, we observed CRISPLD2 has been shown to be immune-responsive, in that patients with asthma, chronic inflammation in the lungs, exhibit significant expression levels of CRISPLD2 following glucocorticoid treatment.^62^ Glucocorticoids (GCs) are anti-inflammatory, highlighting that anti-inflammatory actions in the lung via GCs occurred in conjunction with increases in CRISPLD2 gene expression. Himes et al. investigated this further and found CRISPLD2 to be an immune-responsive gene that increased in response to proinflammatory cytokine IL1ß.^62^ We identified an increase in CRISPLD2 expression in our pre- vs. post-blast exposure gene expression analyses (Figure 3B), highlighting the presence of an inflammatory response following exposure to blast overpressure waves in our study. We further observe inflammation-related gene expression in the downregulation of KCTD12 (Figure 3B), which was shown to be under expressed in a genome-wide expression microarray study by Miller et al. investigating chronic stress within caregivers of brain cancer patients vs. controls^63^, thus indicating a repressive effect of chronic stress on KCTD12 expression.

Additionally, other inflammatory pathways enriched in our gene expression dataset include the NF-kB mediated TNF-alpha signaling gene hallmark set, with the highest normalized enrichment score (Supplementary Table S7). The NF-kB pathway responds to elevation of proinflammatory cytokine TBF-alpha, and further NF-kB activation is known to occur in both acute inflammatory responses and in chronic inflammatory diseases.^64–66^ Acute changes in circulating inflammatory cytokines following blast exposure were also observed in our assessment of serum cytokine levels across multiple timepoints of the breacher operations training course. We observed an increase in MCP-1, GCSF, HGF, MCSF (CSF-1), and RANTES on days of blast exposure (Figure 5, Days 7 and 9). Notably, MCP-1 (monocyte chemoattractant protein-1, CCL-2) has been shown to be involved in neuron-immune cell interactions and more specifically, is responsible for macrophage movement into peripheral nerves, microglia activation, and recently has been implicated as modulated by the transcription factor ATF3 following traumatic brain injury.^67–70^ Generally, these cytokines play a role in the recruitment of white blood cells to injured areas ^71^, which supports previous observations on inflammatory response following blast exposure, even in short (day to day) time frames.^72–74^

Moreover, blast-associated physiological and psychological symptom analyses in participants with low vs. high cumulative blast exposure revealed that tinnitus was the primary symptom present using daily symptom reports throughout the breaching operations course. This finding is not surprising because not only is tinnitus a commonly reported symptom by military and Veteran populations, but it is in fact the most prevalent service-connected disability of all Veterans Benefit Administration compensation recipients, according to the 2018 VA Annual Benefits Report.^75^ DNA methylation and transcriptional profiles were compared in the low vs. high cumulative blast groups, and although we identified no significant genome wide transcriptional changes following multiple testing correction, we did identify genes with differential DNA methylation changes that track with the reported symptom of tinnitus in the low vs. high career blast groups (Figure 4B). Two of the genes, KCNE1 and CYP2E1, showed the expected inversely correlated pattern of increase in DNA methylation and decrease in gene expression. KCNE1 is a potassium channel that has been previously implicated in noise induced hearing loss (NIHL) and tinnitus through genetic studies^43, 44^, and has been shown to play a critical role in K+ recycling in the endolymph of the inner ear.^76^ CYP2E1 metabolizes acrylonitriles, which have been demonstrated to promote NIHL through oxidative stress mechanisms.^45^ We also observed changes in hearing-related genes in our pre-post blast exposure gene expression analysis, in that KCTD12 which was robustly downregulated has been associated with tinnitus.^77^ This is of particular interest to the Veterans Affairs (VA) community because tinnitus is a commonly reported symptom by military and Veteran personnel with repeated exposure to blast. Our knowledge of the neurobiological mechanisms behind auditory and vestibular injuries in our military warfighters is limited but particularly relevant given that tinnitus is present in 11% of OIF and OEF Veterans and has been demonstrated in active duty medical records to be the primary clinical risk for personnel in military occupational specialties (MOSs) associated with blast exposure.^78^ Our results suggest that these hearing-related genes may play a direct role in auditory processing and the perception of sound.

Overall, we demonstrate that blast exposure, both acutely and accumulation, is capable of altering gene expression and DNA methylation patterns, respectively. Further epigenetic, transcriptional, and inflammatory studies are required to deduce the molecular biosignatures of blast and the associated symptoms.

## CONCLUSION

As DNA methylation represents highly stable, long lasting marks, these are likely representative of cumulative blast exposures, whereas changes in gene transcription are facile and represent acute or proximal exposure to blast events. Hence, it is not surprising that we did not detect significant gene expression changes at baseline associated with lifetime blast exposure. We did, however, detect changes in gene expression directly following blast exposure representing a rapid molecular response to blast overpressure blast. The present study suggests that physiological responsivity to different environmental factors, in this case blast overpressure exposure, may be captured by differing biomarkers–with DNA methylation encapsulating the chronic cumulative exposures and inflammatory & RNA transcription the acute response. This systems-based approach allows for context-dependent investigations, allowing for discovery of molecular perturbations and symptomatology both proximally to exposure to blast overpressure and distally across the lifespan.

## ACKNOWLEDGEMENTS

This work was supported by the U.S. Army Medical Research and Development Command/USAMRDC and the Veterans Affairs Office of Research and Development. The opinions or assertions contained herein are the private views of the authors, and are not to be construed as official, or as representing the true views, position or policy of the U.S. Government, the Department of the Army, Department of the Navy, the Defense and Veterans Brain Injury Center, the Department of Defense, or the Department of Veterans Affairs. The investigators have adhered to the policies for protection of human subjects as prescribed in AR 70-25. This research was supported in part by an appointment to the Research Participation Program at the Walter Reed Army Institute of Research administered by the Oak Ridge Institute for Science and Education through an interagency agreement between the U.S. Department of Energy and Medical Research and Development Command/USAMRDC. WRAIR and NMRC support further derives from Joint Program Committee Five intramural awards, The Development of Blast Exposure Standards, Evaluation of the Effects of High Level Overpressure (8+ psi) on Cognitive Performance, Brain Blood Biomarkers and Symptom Reporting, and Environmental Sensors in Training (ESiT). The study protocol was approved by the Walter Reed Army Institute of Research and the Naval Medical Research Center Institutional Review Board’s in compliance with all applicable federal regulations governing the protection of human subjects. Some of the authors are military service members or federal/contracted employees of the United States government. This work was prepared as part of their official duties. Title 17 U.S.C. 105 provides that copyright protection under this title is not available for any work of the United States Government. We would like to thank the instructors and students from the Urban Mobility Breaching Course at site 3 for volunteering to participate in this study; the WRAIR field research team for collecting biological samples and performance data as well as formatting the data for analytical use; and Dr. Angela Boutté for her critical review and editorial comments of the manuscript. Dr. Haghighi’s research is supported by RX001705, CX001395, BX003794, and CX001728.

## CONFLICT OF INTEREST

The authors declare that they have no competing interests.

---

**Figure S1:**
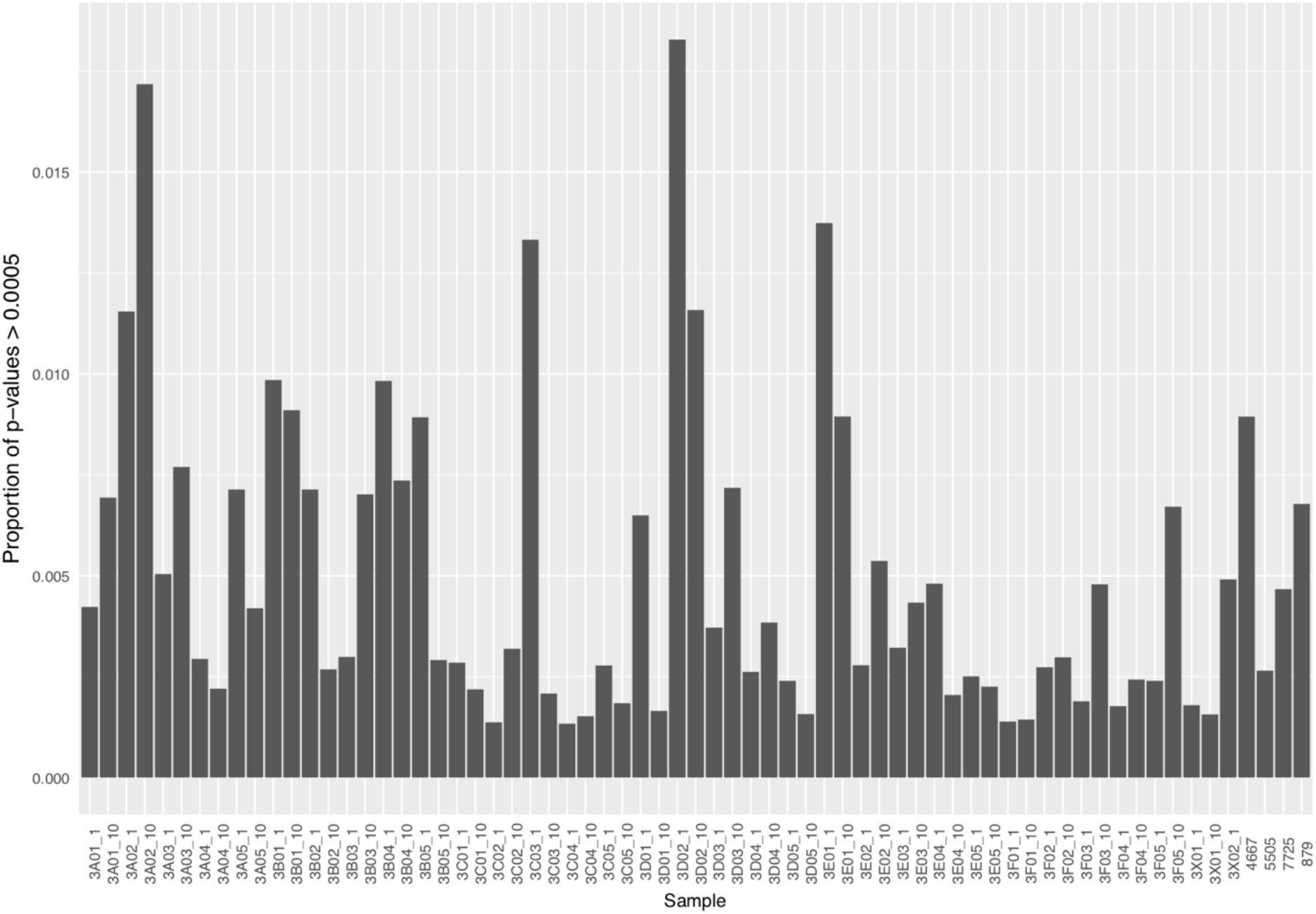
Quality control, detection p-value. Showing the proportion of probes with poor quality (p>0.0005) probes for each biological sample. All samples had high quality data, with at least 98% of probes passing criteria.

**Figure S2:**
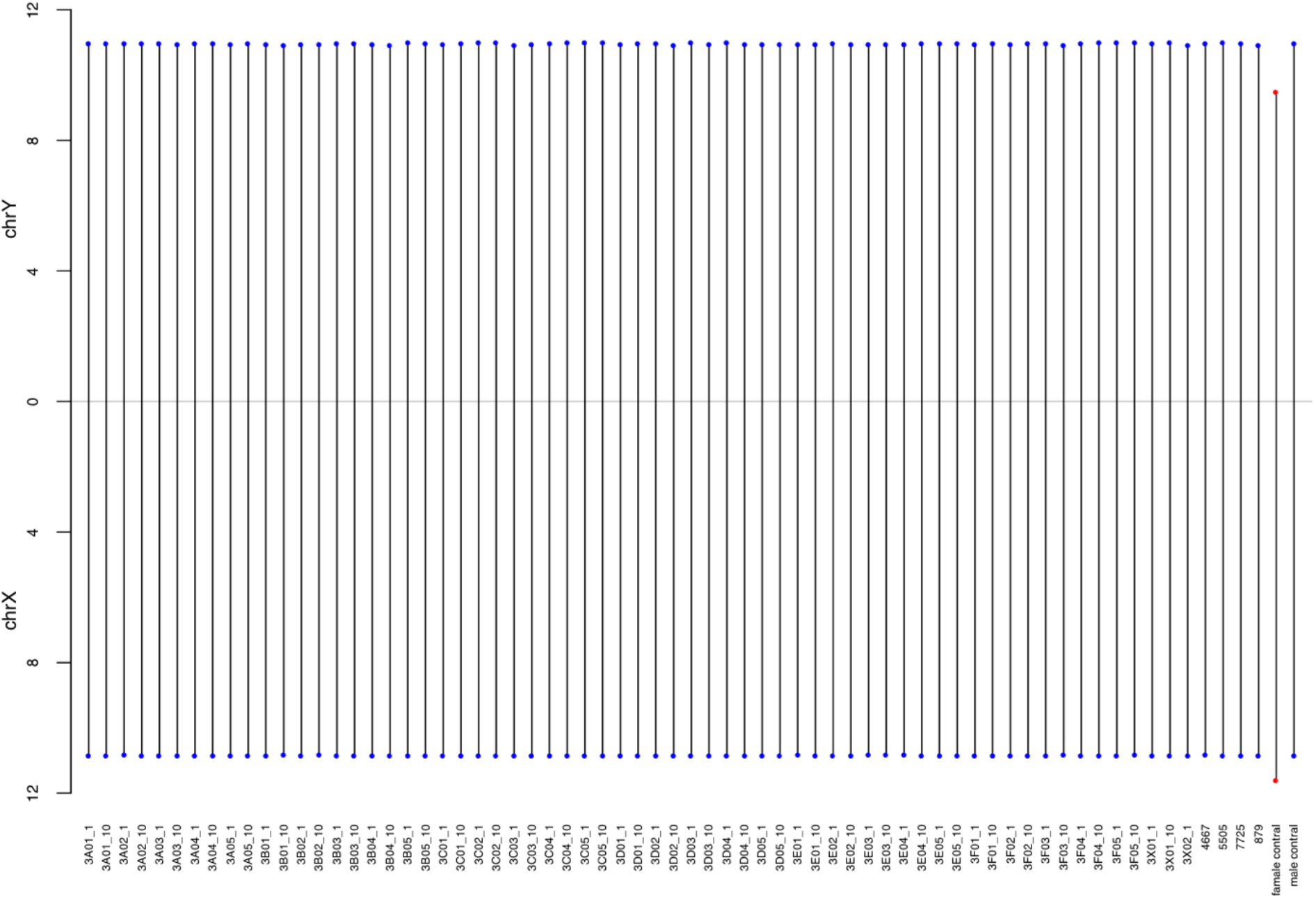
Quality Control, gender prediction. Showing QC plot for gender prediction demonstrating consistency for predicted and reported gender, using Chromosome X and Chromosome Y median intensity. Known male and female reference samples were added as experimental controls. Blue circles are males and red circles are females.

**Figure S3:**
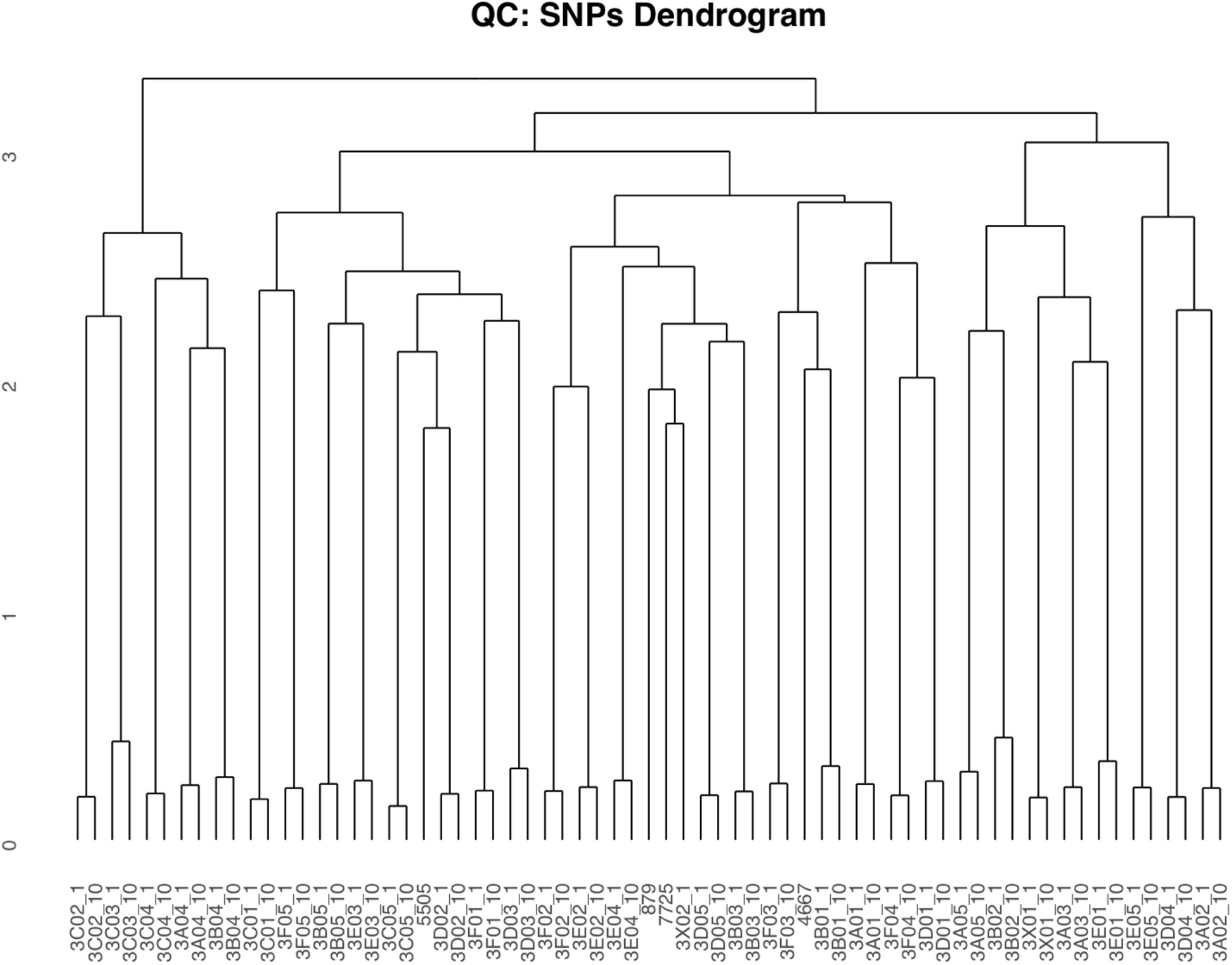
Quality Control, genotype consistency check. QC of paired samples pre- and post-training exposure. Dendrogram showing clustering of individuals by genotypes derived from the 65 SNP probes for biological samples collected pre/post training. The y-axis shows Euclidean distance and samples that show no clustering were those for which no post-training biological samples were available.

**Figure S4:**
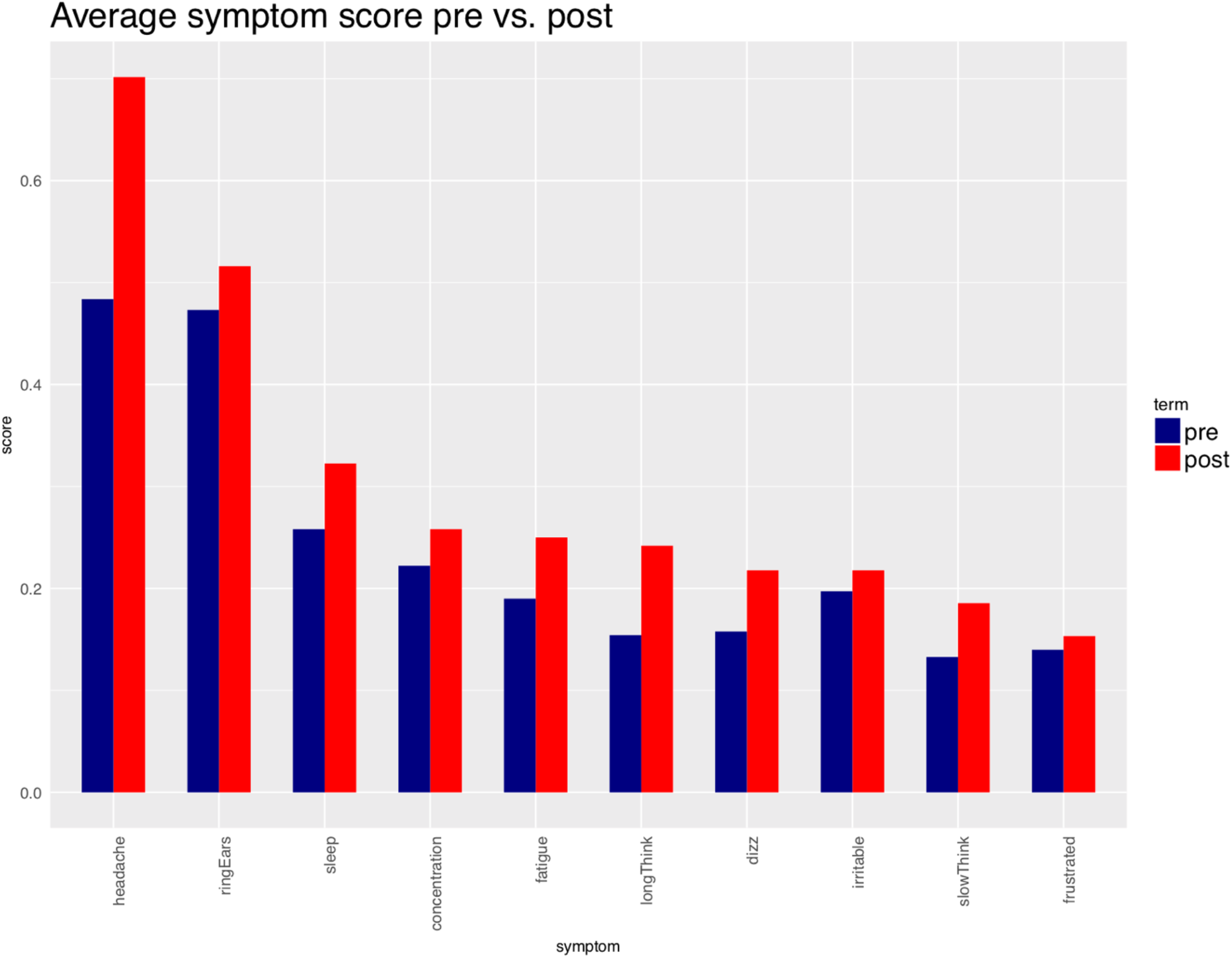
Average symptom score pre-post blast exposure, with headache being the highest reported symptom (endorsed by 18 participants).

**Figure S5:**
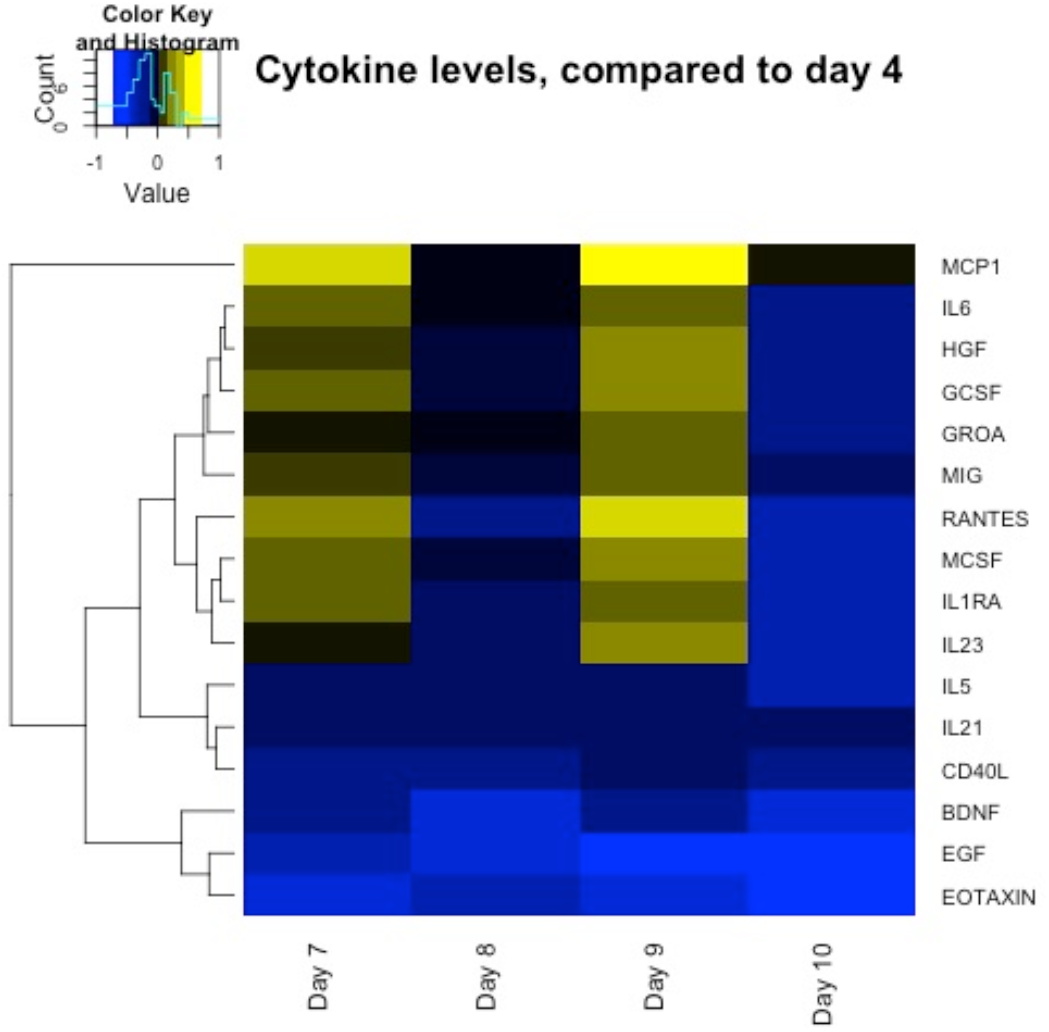
Heatmap showing extended cytokine panel with significant changes in cytokine level tracking with blast exposure (days 7 and 9) using training day 4 as the reference.

